# Excess histone H3 is a Chk1 inhibitor that controls embryonic cell cycle progression

**DOI:** 10.1101/2020.06.09.142414

**Authors:** Yuki Shindo, Amanda A. Amodeo

## Abstract

The early embryos of many species undergo a switch from rapid, reductive cleavage divisions to slower, cell fate-specific division patterns at the Mid-Blastula Transition (MBT). The maternally loaded histone pool is used to measure the increasing ratio of nuclei to cytoplasm (N/C ratio) to control MBT onset, but the molecular mechanism of how histones regulate the cell cycle has remained elusive. Here, we show that excess histone H3 inhibits the DNA damage checkpoint kinase Chk1 to promote cell cycle progression in the *Drosophila* embryo. We find that excess H3-tail that cannot be incorporated into chromatin is sufficient to shorten the embryonic cell cycle and reduce the activity of Chk1 *in vitro* and *in vivo*. Removal of the Chk1 phosphosite in H3 abolishes its ability to regulate the cell cycle. Mathematical modeling quantitatively supports a mechanism where changes in H3 nuclear concentrations over the final cell cycles leading up to the MBT regulate Chk1-dependent cell cycle slowing. We provide a novel mechanism for Chk1 regulation by H3, which is crucial for proper cell cycle remodeling during early embryogenesis.

---

The DNA damage checkpoint is crucial to protect genome integrity^1,2^. However, the early embryos of many species sacrifice this safeguard to allow for rapid cleavage divisions that are required for speedy development. At the Mid-blastula transition (MBT), the cell cycle is remodeled with the addition of gap-phases and acquisition of DNA damage checkpoints, resulting in dramatic lengthening in cell cycle durations. DNA damage checkpoint activity is essential for proper cell cycle slowing at the MBT in *Drosophila*^3–9^. At the same time, major zygotic transcription is activated for the first time. The nuclear-to-cytoplasmic (N/C) ratio determines the timing of both cell cycle slowing and transcription initiation^10–14^ (Fig. 1A). How the N/C ratio is sensed has been an area of intense study, but the prevailing model is that a maternally loaded transcriptional inhibitor or cell cycle activator is titrated against the exponentially increasing number of nuclei^9–11,15,16^. Histones have emerged as a strong candidate for the titrated molecule as the pool size of maternally provided histones is critical for proper MBT timing in frogs, fish, and flies^17–22^. The regulatory activity of histones appears to be upstream of the checkpoint kinase, Chk1^22^, but how histone availability feeds into the cell cycle, whether it is direct or indirect, has remained unclear.

**Fig. 1.**
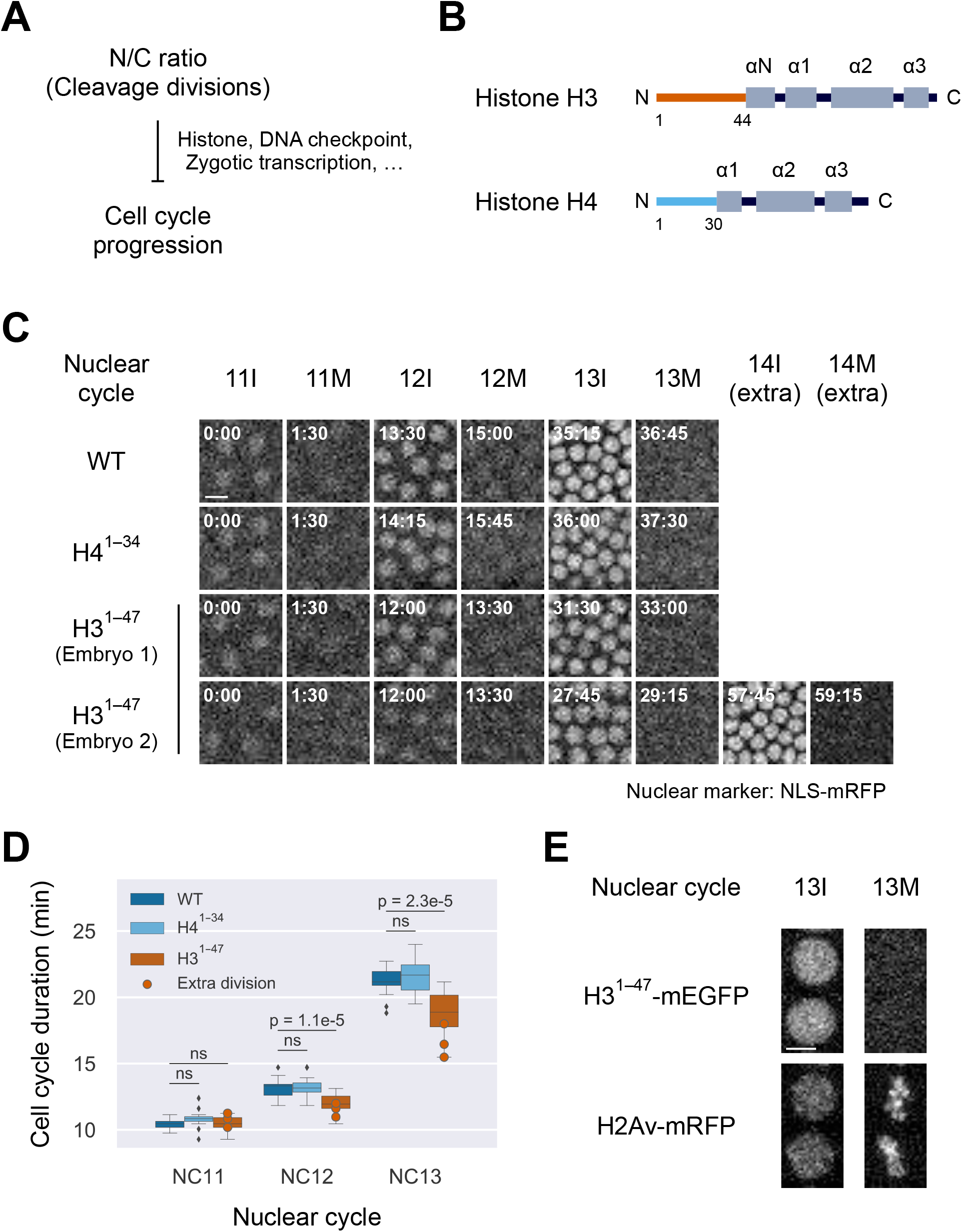
Histone H3 regulates the cell cycle in the early embryo independent of chromatin incorporation. (A) The nuclear to cytoplasmic (N/C) ratio exponentially increases with each cleavage division cycle and inhibits cell cycle progression when a threshold N/C ratio is met in the early embryo. Multiple mechanisms, including histone titration, DNA damage checkpoint, and zygotic transcription have been implicated to contribute to this transition, although the precise molecular mechanism has remained poorly understood. (B) Schematic of the secondary structure of histones H3 and H4, where α-helices are shown as boxes. The structural information is based on HistoneDB 2.0^23^. (C) The N-terminus of histone H3 (H3^1-47^) and histone H4 (H4^1-34^) tagged with mEGFP were each maternally expressed and nuclei were visualized by a NLS-mRFP over the course of syncytial blastoderm stage. Embryos expressing H3^1-47^ had faster cell cycles compared to WT and ~20% underwent an extra division in NC14, indicating that H3^1-47^ interferes with cell cycle slowing. H4-tail did not affect cell cycle duration or number. Scale bar, 10 μm. (D) Cell cycle times as measured by durations between nuclear envelope breakdown are shown as boxplots, where the box represents the 25^th^ and 75^th^ percentiles, the bar in the middle of the box represents the median, the whiskers extend to points that lie within 1.5 IQRs of the box, and points that fall outside the whiskers are shown by diamonds. Embryos that underwent an extra division at NC14 are also plotted by circles. WT embryos always stop in NC14, while H3^1-47^-mEGFP embryos frequently undergo one extra division. ns, not significant. (E) Confocal images of H3^1-47^-mEGFP and H2Av-mRFP during a nuclear division at NC13. H3^1-47^-mEGFP is imported into the nucleus during interphase but not incorporated into chromatin and lost in mitosis. Scale bar, 5 μm.

Given the well-established function of histones as components of chromatin, previous work has focused on the role of histone abundance in regulating chromatin state and zygotic genome activation^17–22^. To test whether chromatin incorporation was essential for histone regulation of the MBT, we expressed truncated H3 (H3^1-47^) and H4 (H4^1-34^) in the early *Drosophila* embryo. These constructs retain the unstructured tail regions which are the sites of the majority of histone post-translational modifications but lack the histone fold domains required for chromatin incorporation^23^ (Fig. 1B) (hereafter referred to as H3-tail and H4-tail, respectively). We focused on histones H3 and H4 because depletion of all of the replication-coupled histones resulted in a severe defect in the early embryonic cell cycle while a reduction in H2A, H2B, and H2Av had little effect on the early embryogenesis in *Drosophila*^22,24^. Moreover, H3 depletion alone is sufficient for premature MBT in *Xenopus*^20^. We maternally expressed the histone tails and measured cell cycle durations over the course of the syncytial blastoderm stage (Fig. S1). We found that H3-tail expression was sufficient to reduce cell cycle slowing, resulting in embryos with significantly faster cell cycles than WT and H4-tail embryos (Fig. 1, C and D). Additionally, ~20% (3/16) of the H3-tail expressing embryos underwent one extra division prior to cellularization (Fig. 1C and Table S1). Since H3-tail is not incorporated into nucleosomes (Fig. 1E), we concluded that histone H3 can regulate the cell cycle without chromatin incorporation and therefore any changes to the chromatin structure that result from changing histone availability are not essential for the effects of H3 on the cell cycle.

It is well established that in *Drosophila* cell cycle slowing at the MBT requires the activation of Chk1 and regulation of its downstream targets^3–9,25–28^ (Fig. 2A). We had previously observed that total histone reduction resulted in greater activation of Chk1 and corresponding lengthening in cell cycle durations in the early embryo^22^. Therefore, we asked if the H3-tail similarly affected Chk1 activity using a previously described Chk1 sensor^8^. We found that Chk1 activity was decreased in the presence of excess H3-tail in the final two cell cycles before the MBT (Fig. 2B). Therefore, H3-tail acts as an upstream inhibitor of Chk1 activity in the early embryo.

**Fig. 2.**
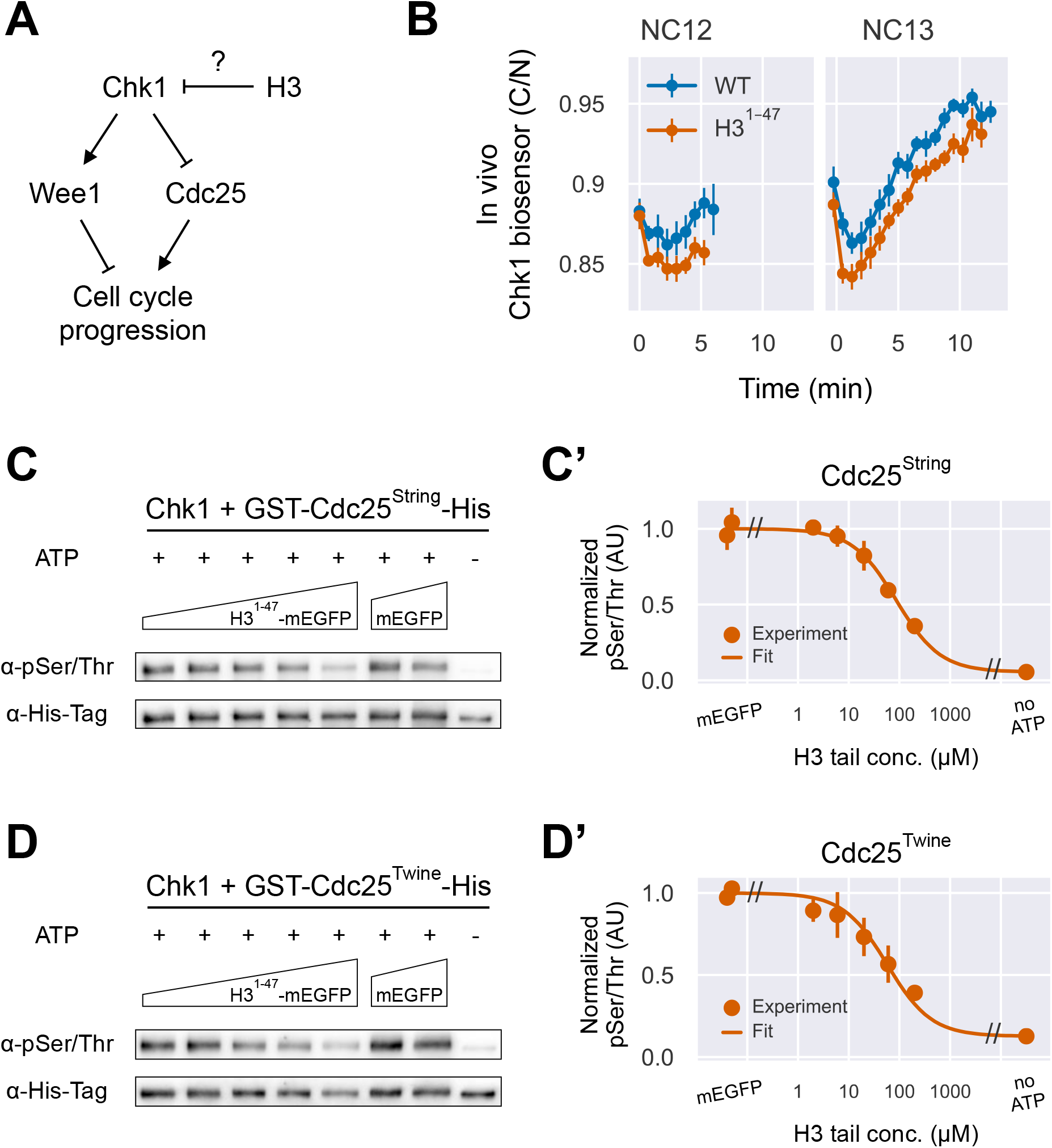
Excess H3-tail inhibits Chk1 activity *in vivo* and *in vitro*. (A) Checkpoint kinase 1 (Chk1) becomes activated at the onset of the MZT and results in cell cycle slowing. This is mediated by multiple Chk1 targets, including Wee1 and Cdc25 whose activities converge on cyclin-Cdk1 to control cell cycle progression. We hypothesized that excess H3 may inhibit Chk1 activity in the early embryo. (B) *In vivo* Chk1 activity as measured by the cytoplasmic-to-nuclear intensity ratio (C/N) of a Chk1 biosensor in WT (blue) and H3-tail (orange) embryos. Chk1 activity is decreased in the presence of excess H3-tail, indicating an inhibitory role of H3-tail on Chk1 in the early embryo. Data represent the mean ± SEM. (C) and (D) We incubated 0.006 μM of Chk1 and 0.2 μM of either GST-Cdc25^String^-His (C) or GST-Cdc25^Twine^-His (D) with varying concentrations of mEGFP (20 and 200 μM) or H3^1-47^-mEGFP (2–200 μM) and detected phosphorylated Cdc25 and total Cdc25 by western blotting. Excess H3-tail inhibits Chk1 phosphorylation of both Cdc25 isoforms *in vitro*. (C’) and (D’) Levels of phosphorylated Cdc25 as assayed in (C) and (D) are normalized to total Cdc25 (His-Tag) and plotted as a function of H3^1-47^-mEGFP concentrations. Data without ATP and mEGFP are also shown. Data represent the mean ± SD of three independent experiments. Data were fitted with a model describing competitive inhibition (see Materials and Methods).

Next, we sought to understand if H3-tail regulation of Chk1 activity is direct or indirect. Previous work has shown that H3 can directly interact with Chk1 as a substrate^29^. Given that the early embryo is provisioned with unusually large amounts of histones, H3 could potentially act as a competitive inhibitor for other, cell cycle regulatory, Chk1 substrates such as Cdc25^String^ and Cdc25^Twine^. To test if H3-tail can directly inhibit Chk1 activity, we performed *in vitro* kinase assays using recombinant human Chk1 with Cdc25 in the presence of different H3-tail concentrations. We found that Cdc25 phosphorylation is reduced by the presence of H3-tail (Fig. 2, C and D). We observed inhibition of Chk1 by the H3-tail in both of the two *Drosophila* Cdc25 isoforms, Cdc25^String^ and Cdc25^Twine^, with Cdc25^Twine^ showing slightly greater sensitivity to inhibition by H3-tail than Cdc25^String^. Using the dose-response relationship obtained from the *in vitro* kinase assay, we estimated the value of *K_i_*, an inhibition constant for the H3-tail, to be 47 μM. This value is comparable to the nuclear H3 concentrations in the early *Drosophila* embryo obtained by other methods (Fig. S2, see Materials and Methods), suggesting that H3 could indeed inhibit Chk1 at physiological concentrations.

To test if the interaction between H3 and Chk1 plays a role *in vivo*, we introduced a mutation at the known Chk1 phosphorylation site, Thr 11, in the H3-tail (H3^1-47, T11A^)^29^. The T11A mutation prevents Chk1-dependent phosphorylation of Thr 11 (Fig. S3) and resulted in a weaker inhibitor activity *in vitro* (Fig. 3A). Therefore, if the main effect of the H3-tail on the cell cycle was through interaction with Chk1, the mutant H3-tail should not shorten the cell cycle like the wild-type H3-tail *in vivo*. We observed that this single point mutation in the H3-tail abolished its ability to affect cell cycle dynamics in the early embryo (Fig. 3B and Fig. S4). Indeed, the H3^1-47, T11A^ mutant tail embryos had cell cycle times that were similar to embryos that did not express any H3-tail (Fig 3B). By contrast, mutations in phosphosites for Aurora-B kinase at S10 and S28 (S10A/S28A) did not affect the H3-tail’s ability to regulate cell cycle slowing in a similar way to the wild-type H3-tail (Fig. 3B). These results indicate that the H3T11 Chk1 phosphosite plays a significant role in the cell cycle slowing.

**Fig. 3.**
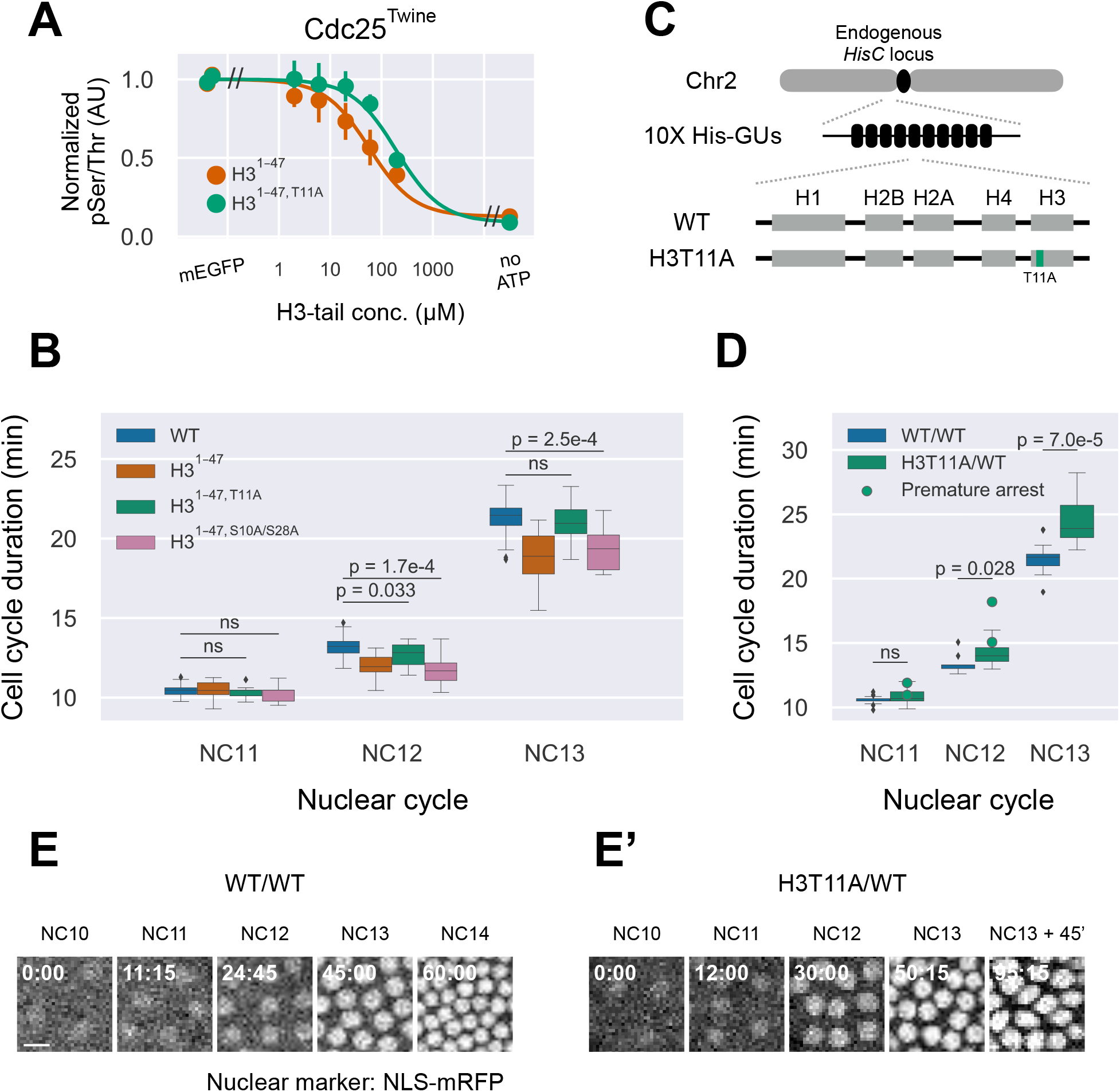
H3 Thr 11 Chk1 phosphosite is necessary for the cell cycle slowing. (A) T11A mutation in H3-tail results in reduced inhibitor activity for Chk1. *In vitro* kinase assays were performed as in Fig. 2D except H3^1-47, T11A^-mEGFP (green) was used in place of H3^1-47^-mEGFP (orange, same as Fig. 2F). Data represent the mean ± SD of three independent experiments. Data were fitted with a model for competitive inhibition. The *K_i_* for H3^1-47, T11A^-mEGFP was 159 μM, 3.4 times larger than that of H3^1-47^-mEGFP (47 μM). (B) Cell cycle times in embryos expressing H3-tail with a T11A mutation or S10A/S28A double mutations. H3-tail with T11A Chk1 phosphosite mutation restored the normal cell cycle while S10A/S28A Aurora B kinase phosphosite mutations resulted in fast cell cycle times similar to wild-type H3-tail. The box represents the 25th and 75th percentiles, the bar in the middle of the box represents the median, the whiskers extend to points that lie within 1.5 IQRs of the box, and points that fall outside the whiskers are shown by diamonds. ns, not significant. (C) Schematic of the engineered histone gene cluster^30^. The engineered gene cluster consists of 10 repeats of a *HisC* gene unit (10X His-GUs) that encodes each of the replication-coupled histones with or without H3T11A mutation. The endogenous *HisC* cluster was replaced by the 10X His-GUs at the endogenous locus. (D) Cell cycle times in the WT His-GUs (10X His-GUs^WT^/10X His-GUs^WT^) and heterozygous H3T11A (10X His-GUs^H3T11A^/10X His-GUs^WT^) embryos are shown as boxplots. Circles indicate embryos that ceased cleavage divisions one cell cycle prematurely. ns, not significant. (E) Confocal images of NLS-mRFP in a WT His-GUs embryo from NC10 through NC14. (E’) Same as (E) except the embryo was heterozygous for H3T11A, in which nuclei stopped dividing one cell cycle prematurely.

To test the role of the Chk1 phosphosite in the context of the endogenous H3, we took advantage of a recently-developed genetic system^30^ in which all of the endogenous replication-coupled histone genes have been replaced by an artificial histone gene cluster and each of the known histone modification sites have been individually mutated (Fig. 3C). Using the H3T11A mutant from this collection, we analyzed embryos produced from heterozygous H3T11A females, since homozygotes are sterile. Note that the single amino acid mutation in the histone gene cluster did not affect total histone levels (Fig. S5). We found that cell cycle durations in the heterozygous H3T11A embryos lengthened compared to WT, indicating that the decrease in the amount of Chk1-phosphorylatable H3 promoted cell cycle slowing (Fig. 3D). Additionally, ~15% (2/14) of the heterozygous H3T11A embryos ceased cleavage cycles one cell cycle prematurely (Fig. 3E, Fig. S6, and Table S1). Together, the complementary behaviors of the cell cycle phenotype in response to either overexpression or depletion of H3T11 phosphosite indicates that the amount of H3T11 times DNA checkpoint activation that controls the cell cycle progression during embryogenesis.

Given the experimental observations described above, we propose that histone H3 regulates cell cycle progression through Chk1 inhibition in the early embryo. Since nuclear concentration of H3 decreases as the N/C ratio increases^31^, this mechanism could provide an explanation of how the early embryo couples the change in the number of nuclei to cell cycle slowing and developmental progression. To quantitatively test this idea, we constructed a computational model of the cyclin-Cdk1 system that drives the embryonic cell cycle^32,33^ and introduced Chk1 into the model (Fig. 4A). When Chk1 activity is relatively low, the model exhibited stable oscillations, while increased Chk1 activity resulted in a loss of the oscillatory behavior and the system converged to a single stable fixed point, which can be interpreted as cell cycle arrest (Fig. 4B), consistent with the observation that Chk1 activity is necessary for proper cell cycle slowing^3–9^. In our model, the apparent cyclin synthesis rate was decreased slightly with each nuclear cycle to account for the small increase in cell cycle duration that persists in Chk1 mutant embryos^5,7^ (Fig. S7). We next modeled Chk1 activity as dependent on H3 concentration using experimentally constrained parameter values and asked how H3 availability affects cell cycle dynamics. We found that the experimentally measured decrease in nuclear H3 over the course of the final cleavage division cycles well recapitulates the dynamics of cell cycle slowing at the MBT through decreasing Chk1 inhibition (Fig. 4C). The model quantitatively recapitulated cell cycle durations of H3-tail expression (Fig. 1C), H3T11A-tail expression (Fig. 3B), heterozygous H3T11A replacement (Fig. 3D), and H3 depletion^22^ (Fig. 4D). We also observed premature cell cycle arrest in NC13 when we depleted 70% of H3, slightly more than 60% that we previously experimentally determined in bulk^22^, suggesting that the bifurcation of the system occurs in a realistic range of H3 depletion (Fig. S8). Given that embryo-embryo variabilities may exist in the living embryos, the model is consistent with the previous observation that approximately half of the H3 depleted embryos underwent either partial or full cell cycle arrest in NC13^22^. Together, our computational model quantitatively supports the idea that Chk1 inhibition by H3 allows the early embryo to couple the cleavage division cycles to the regulation of cell cycle remodeling at the MBT.

**Fig 4.**
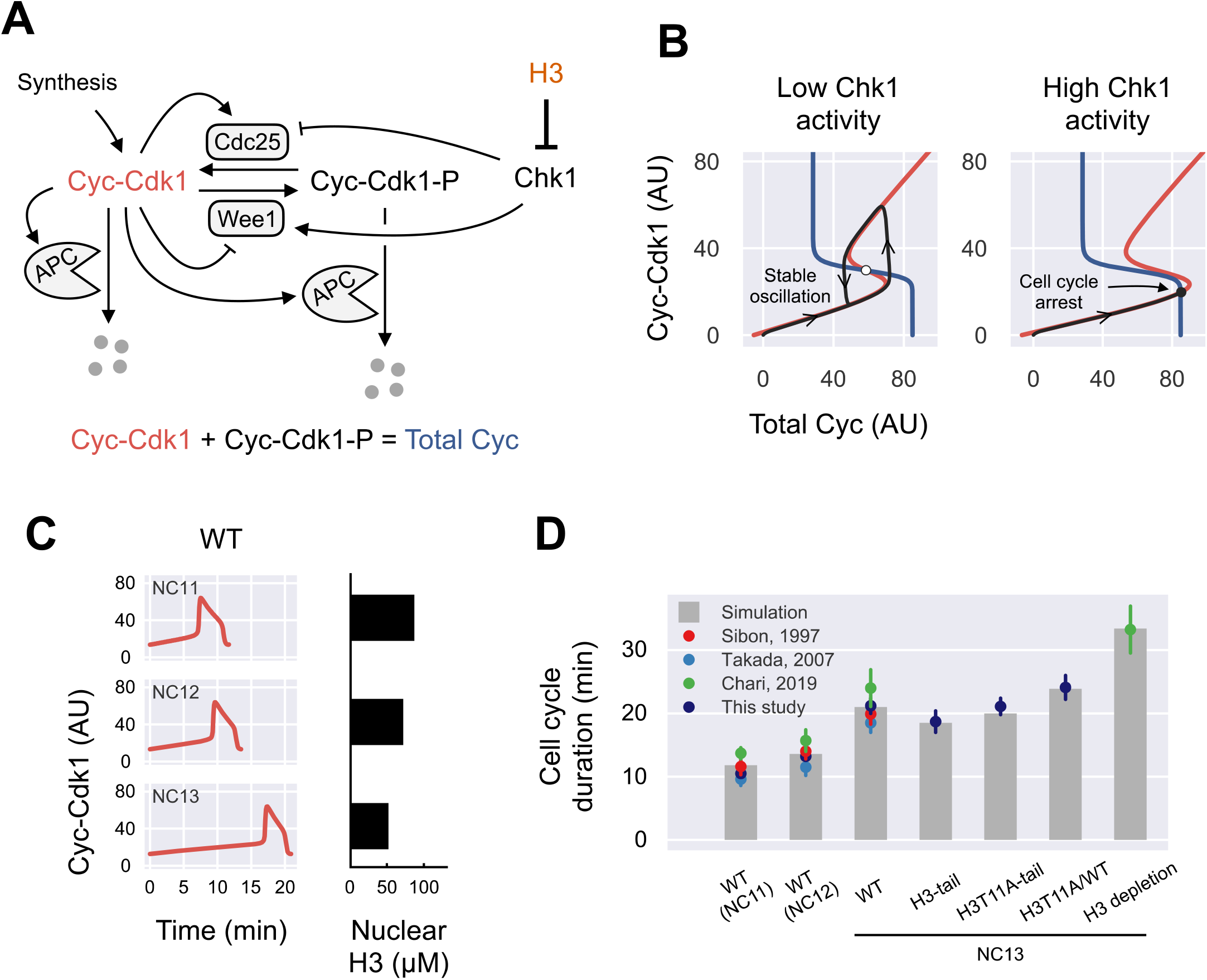
A mathematical model for N/C ratio sensing through Chk1 inhibition by H3 recapitulates cell cycle phenotypes. (A) Schematic of the model. (B) Phase planes of the model with respect to low (left) and high (right) Chk1 activity. The total Cyc and Cyc-Cdk1 nullclines (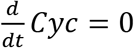 and 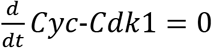) are shown in blue and red, respectively. Black lines represent sample trajectories from an initial condition of (0, 0). When Chk1 activity is relatively low (left), the intersection of the nullclines is unstable (open circle) and all trajectories converge to a stable limit cycle. In the presence of high Chk1 activity (right), the nullclines intersect at a stable steady state (filled circle) that corresponds to cell cycle arrest. (C) Simulation of Cyc-Cdk1 oscillations in NC11–13 in the WT embryo. Titration of nuclear H3 by the increasing number of nuclei (Fig. S2) resulted in lengthening in cell cycle durations. (D) Simulated cell cycle durations are shown along with experimental data obtained by previous studies and this study. The model quantitatively recapitulates the cell cycle dynamics, including cell cycle slowing in WT and changes in NC13 durations in response to various perturbations.

The N/C ratio has long been known to control the switch from synchronous rapid divisions to slower, cell fate-specific division patterns at the MBT, but how it is sensed has remained unclear. Here, we propose that histone H3 is used to sense the N/C ratio through direct competitive inhibition of Chk1. We have shown that most nuclear H3 is non-DNA-bound in the early cleavage division cycles, but the free pool becomes depleted by the exponentially increasing N/C ratio in final pre-MBT cycles, resulting in the decrease in nuclear H3 concentrations with each cycle^31^. We propose that the excess maternally provided H3 inhibits Chk1 to prolong rapid cleavage cycles until a sufficient N/C ratio threshold is met. Chk1 inhibition in early cycles is consistent with the observation that a DNA damage response is suppressed in pre-MBT embryos and that addition of sufficient exogenous DNA triggers premature cell cycle slowing *in vivo* and checkpoint activity *in vitro*^34,35^. Our model not only provides a simple molecular mechanism to couple cell cycle slowing and developmental progression to cell number and cell size, it also demonstrates a novel role for unincorporated, free histones as regulatory molecules.

## Acknowledgments

We thank Allan C. Spradling, Robert J. Duronio, and Yasushi Sako for plasmids; Stefano Di Talia, Junbiao Dai, and the Bloomington *Drosophila* Stock Center (NIH P40OD018537) for fly stocks; Jan Skotheim, Kurt Schmoller, Stanislav Shvartsman, Trudi Schüpbach and Eric Wieschaus for discussion and comments on the manuscript. We also thank Gary Laevsky and the Confocal Imaging Facility at Princeton University and Gordon Gray and the *Drosophila* Media Core Facility at Princeton University. Y.S. is supported by JSPS Overseas Research Fellowships, the Postdoctoral Fellowship from the Uehara Memorial Foundation, and the Osamu Hayaishi Memorial Scholarship for Study Abroad.

## Author contributions

Conceptualization, Y.S and A.A.A.; Investigation, Y.S. and A.A.A; Writing – Original Draft, Y.S. and A.A.A.; Writing – Review & Editing, Y.S. and A.A.A.

