## Supplemental MM and Figs for "Excess histone H3 is a Chk1 inhibitor that controls embryonic cell cycle progression"

### Methods

#### Reagents

Reagents and fly stocks we used in this study are summarized below.

| Reagent or Resource | Source | Identifier |
|---|---|---|
| Anti-H2B | Abcam | ab52484 |
| Anti-H3 | Abcam | ab1791 |
| Anti-phosphoserine/threonine | ECM Biosciences | PP2551 |
| Anti-His-Tag | Cell Signaling Technology | 2366 |
| Anti-phospho Histone H3 (T11) | Abcam | ab5168 |
| Anti-GFP | Millipore-Sigma | G1546 |
| Alexa Fluor 488 goat anti-mouse IgG | Invitrogen | A-11001 |
| Alexa Fluor 546 goat anti-mouse IgG | Invitrogen | A-11003 |
| Alexa Fluor 647 goat anti-rabbit IgG | Invitrogen | A-21244 |
| Rosetta 2 Competent Cells | Millipore-Sigma | 71402-3 |
| Protease Inhibitor Cocktail | Millipore-Sigma | P8465 |
| cOmplete His-tag Purification Resin | Millipore-Sigma | 5893682001 |
| Imidazole | Millipore-Sigma | I202 |
| Glutathione Sepharose 4B | GE Healthcare | 17075601 |
| Glutathione, Reduced, Free Acid | Millipore-Sigma | 3541 |
| PreScission Protease | GenScript | Z02799 |
| Benzonase Nuclease | Millipore-Sigma | E8263-5KU |
| CHK1, Active | SignalChem | C47-10G |
| yw | Bloomington Stock Center | 1495 |
| yw ; ; UASz-H3$^{1-47}$-mEGFP | This study | N/A |
| yw ; ; UASz-H4$^{1-34}$-mEGFP | This study | N/A |
| yw ; ; UASz-H3$^{1-47}$-miRFP670nano | This study | N/A |
| yw ; ; UASz-H3$^{1-47,\ T11A}$-mEGFP | This study | N/A |
| yw ; ; UASz-H3$^{1-47,\ S10A/S28A}$-mEGFP | This study | N/A |
| NLS-mRFP, w | Bloomington Stock Center | 31418 |
| w ; His2Av-mRFP | Bloomington Stock Center | 23651 |
| w ; matα4-tubGal4 | Bloomington Stock Center | 7062 |
| yw ; Sp / CyO ; Cdc25C$^{183-251}$-EGFP | Di Talia lab | N/A |
| w ; 10x His-GUs$^{WT}$ | Dai lab | N/A |
| w ; 10x His-GUs$^{H3T11A}$ / CyO, ActGFP | Dai lab | N/A |

#### Stocks and crosses

All fly stocks were maintained at room temperature of approximately 22 °C on a standard cornmeal media. Crosses of UASz females or control yw females to matα4-tubGal4 males were carried out at 25 °C.

### Plasmids and transgenesis

The pUASz plasmid was a gift from Allan C. Spradling[1], the pMBAC-1xHisC plasmid was a gift from Robert J. Duronio[2], the GST-mEGFP-ERK2-8XHis plasmid was a gift from Yasushi Sako[3]. Fragments of $H3^{1-47}$, $H4^{1-34}$, and mEGFP were amplified by PCR and assembled into the pUASz plasmid (pUASz-$H3^{1-47}$-mEGFP and pUASz-$H4^{1-34}$-mEGFP). To generate pUASz-$H3^{1-47}$-miRFP670nano, the mEGFP sequence in the pUASz-$H3^{1-47}$-mEGFP plasmid was replaced by a gBlocks Gene Fragments (IDT) encoding miRFP670nano[4]. To generate pUASz-$H3^{1-47,\ T11A}$-mEGFP and pUASz-$H3^{1-47,\ S10A/S28A}$-mEGFP plasmids, the wild-type $H3^{1-47}$ sequence in the pUASz-$H3^{1-47}$-mEGFP plasmid was replaced by a gBlocks Gene Fragment (IDT) encoding either $H3^{1-47,\ T11A}$ or $H3^{1-47,\ S10A/S28A}$. The transgenes were inserted into the VK33 *attP* site on chromosome 3L[5] via phiC31-mediated integration (BestGene). Coding regions of $cdc25^{string}$ and $cdc25^{twine}$ genes were amplified by PCR from yw adult genomic DNA and assembled into the GST plasmid backbone (GST-$Cdc25^{String}$-His, GST-$Cdc25^{Twine}$-His). $H3^{1-47}$-mEGFP, $H3^{1-47,\ T11A}$-mEGFP, and mEGFP sequences were amplified by PCR and inserted into the GST plasmid backbone without C-terminal 8XHis and N-terminal 8XHis tag was then added by PCR for two-step purification (His-GST-$H3^{1-47}$-mEGFP, His-GST-$H3^{1-47,\ T11A}$-mEGFP, and His-GST-mEGFP).

### Genotypes

Females with the genotype described below were mated to yw males and the resultant embryos were analyzed.

Fig. 1. C and D:
WT: NLS-mRFP / y,w ; matα4-tubGal4 / + ; + / +
$H4^{1-34}$: NLS-mRFP / y,w ; matα4-tubGal4 / + ; UASz-$H4^{1-34}$-mEGFP / +
$H3^{1-47}$: NLS-mRFP / y,w ; matα4-tubGal4 / + ; UASz-$H3^{1-47}$-mEGFP / +
The numbers of embryos analyzed are 16, 11, and 15, respectively.

Fig. 1E:
w / y,w ; matα4-tubGal4 / H2Av-mRFP ; UASz-$H3^{1-47}$-mEGFP / +

Fig. 2B:
WT: NLS-mRFP / y,w ; matα4-tubGal4 / + ; $Cdc25C^{183-251}$-EGFP / +
$H3^{1-47}$: NLS-mRFP / y,w ; matα4-tubGal4 / + ; $Cdc25C^{183-251}$-EGFP / UASz-$H3^{1-47}$-miRFP670nano
The numbers of embryos analyzed are 8 and 13, respectively.

Fig. 3B:
WT: NLS-mRFP / y,w ; matα4-tubGal4 / + ; + / +
$H3^{1-47,\ T11A}$: NLS-mRFP / y,w ; matα4-tubGal4 / + ; UASz-$H3^{1-47,\ T11}$A-mEGFP / +
$H3^{1-47,S10A/S28A}$: NLS-mRFP / y,w ; matα4-tubGal4 / + ; UASz-$H3^{1-47,\ S10A/S28A}$-mEGFP / +
The numbers of embryos analyzed are 26, 13, and 14, respectively.

Fig. 3D:

WT/WT: NLS-mRFP / w ; 10X His-GUs$^{WT}$ / 10X His-GUs$^{WT}$
H3T11A/WT: NLS-mRFP / w ; 10X His-GUs$^{H3T11A}$ / 10X His-GUs$^{WT}$
The numbers of embryos analyzed are 15-17, and 10-14, respectively.

Fig. S1:
yw: yw ; + / + ; + / +
H3$^{1\text{-}47}$: NLS-mRFP / y,w ; matα4-tubGal4 / + ; UASz-H3$^{1\text{-}47}$-mEGFP / +

Fig. S4:
H3$^{1\text{-}47}$: NLS-mRFP / y,w ; matα4-tubGal4 / + ; UASz-H3$^{1\text{-}47}$-mEGFP / +
H3$^{1\text{-}47,\ T11A}$: NLS-mRFP / y,w ; matα4-tubGal4 / + ; UASz-H3$^{1\text{-}47,\ T11}$A-mEGFP / +
H3$^{1\text{-}47,S10A/S28A}$: NLS-mRFP / y,w ; matα4-tubGal4 / + ; UASz-H3$^{1\text{-}47,\ S10A/S28A}$-mEGFP / +

Fig. S5:
WT/WT: NLS-mRFP / w ; 10X His-GUs$^{WT}$ / 10X His-GUs$^{WT}$
H3T11A/WT: NLS-mRFP / w ; 10X His-GUs$^{H3T11A}$ / 10X His-GUs$^{WT}$

Fig. S6:
WT/WT: NLS-mRFP / w ; 10X His-GUs$^{WT}$ / 10X His-GUs$^{WT}$
H3T11A/WT: NLS-mRFP / w ; 10X His-GUs$^{H3T11A}$ / 10X His-GUs$^{WT}$

### Microscopy

Dechorionated embryos were mounted in deionized water on a glass-bottom microwell dish (MatTek) and fluorescent images were acquired by a Nikon A1R laser scanning confocal microscope with a 20X 0.75 NA objective at room temperature of approximately 22°C. Time-lapse movies were obtained at a time resolution of 45 s. Cell cycle time was measured by the number of frames between the nuclear envelope breakdown in a given cell cycle and in the next cell cycle. To correct for day-to-day temperature fluctuations in our microscope facility that could affect the cell cycle measurements, we imaged both WT and genetic manipulation of interest in the same dish at the same time and cell cycle times were normalized to the average duration of WT NC11[6]. For measurements of *in vivo* Chk1 activity[7], nuclear regions were segmented from NLS-mRFP images using ilastik[8], eroded by one pixel to ensure that only the nuclear signal was measured, and nuclear Cdc25C$^{183\text{-}251}$-EGFP intensities were quantified. Cytoplasmic masks were generated by 3-pixel-dilation of the nuclear masks followed by subtraction of 1-pixel-dilated nuclear masks to ensure that only the cytoplasmic signal is measured, and cytoplasmic intensities of the Chk1 biosensor were quantified.

### Purification of recombinant proteins

The plasmids were transformed into *E. coli* Rosetta 2, and recombinant protein expression was induced for 16 h at 18 °C by the addition of 0.1 mM IPTG. For His-GST-mEGFP, His-GST-H3$^{1\text{-}47}$-mEGFP, His-GST-H3$^{1\text{-}47,\ T11A}$-mEGFP, His-GST-H3$^{1\text{-}47,\ S10A/S28A}$-mEGFP, and GST-Cdc25$^{Twine}$-His proteins, lysates were recovered in 20 mM Tris-Cl (pH 7.5), 150 mM NaCl, 1 mM

DTT, and 1X Protease Inhibitor Cocktail (Sigma), incubated with cOmplete His-tag Purification Resin, and eluted in 20 mM Tris-Cl (pH 7.5), 150 mM NaCl, and 250 mM Imidazole. Benzonase Nuclease and $MgCl_2$ were added to final concentrations of 250 units/ml and 1 mM, respectively, and the samples were incubated overnight at 4 °C to remove any contamination of nucleic acids. Then the samples were incubated with Glutathione Sepharose 4B. For mEGFP, $H3^{1-47}$-mEGFP, and $H3^{1-47,\ T11A}$-mEGFP, the N-terminal His-GST tag was cleaved by the addition of PreScission Protease (GenScript) in 20 mM Tris-Cl (pH 7.5), 150 mM NaCl, 1 mM EDTA, and 1 mM DTT for overnight and eluted proteins were concentrated using Amicon Ultra-4 30 kDa Centrifugal Filter Units. GST-$Cdc25^{Twine}$-His was eluted in 20 mM Tris-Cl (pH 8.0) and 10 mM reduced glutathione and dialyzed twice against 20 mM Tris-Cl (pH 7.5), 150 mM NaCl, 1 mM EDTA, and 1 mM DTT. Since GST-$Cdc25^{String}$-His formed inclusion bodies, purification of GST-$Cdc25^{String}$-His was performed in a denaturing condition. To recover the GST-$Cdc25^{String}$-His in a soluble fraction, the pellet was washed twice in 4% Triton X-100 and twice in sterilized MilliQ water and incubated in 20 mM Tris-Cl (pH 8.0), 300 mM NaCl, 8 M urea, and 10 mM DTT for 1 hour at 37 °C. After centrifugation for 20 min at 4 °C at 13,000 rpm, the supernatant containing GST-$Cdc25^{String}$-His was recovered. The sample was incubated with cOmplete His-tag Purification Resin, eluted in 20 mM Tris-Cl (pH 8.0), 300 mM NaCl, 8 M urea, and 250 mM Imidazole, and dialyzed twice against 20 mM Tris-Cl (pH 8.0), 300 mM NaCl, 1 mM EDTA, and 1 mM DTT. The purified proteins were aliquoted, flash-frozen, and stored at -80 °C until use.

### ***In vitro*** **kinase assay**

For the *in vitro* kinase assay of Chk1 and H3-tail, we incubated 0.06 μM of Chk1 and 2 μM of mEGFP, $H3^{1-47}$-mEGFP, or $H3^{1-47,\ T11A}$-mEGFP in 50 mM Tris-Cl (pH 7.5), 10 mM $MgCl_2$, 2 mM DTT, and 0.25 mM ATP at 25 °C for 15 min in a Bio-Rad T100 thermal cycler. For the *in vitro* kinase assay of Chk1 and Cdc25, we incubated 0.006 μM of Chk1 and 0.2 μM of GST-$Cdc25^{String}$-His or GST-$Cdc25^{Twine}$-His with mEGFP (20 and 200 μM), $H3^{1-47}$-mEGFP (2, 6, 20, 60, and 200 μM), or $H3^{1-47,\ T11A}$-mEGFP (2, 6, 20, 60, and 200 μM) in 50 mM Tris-Cl (pH 7.5), 10 mM $MgCl_2$, 2 mM DTT, and 0.25 mM ATP at 25 °C for 15 min. The reaction was stopped by the addition of 2X Laemmli's sample buffer and boiling at 98 °C for 5 min. Phosphorylated H3T11 and total mEGFP-tagged proteins were detected by western blotting using anti-phospho Histone H3 (T11) and anti-GFP antibodies. Levels of phosphorylated Cdc25 and total Cdc25 proteins were detected by western blotting using anti-phosphoserine/threonine and anti-His-Tag antibodies.

To analyze the effect of H3-tail on Chk1 phosphorylation of Cdc25, levels of phosphorylated Cdc25 were normalized over total Cdc25 levels and then rescaled against control samples in which mEGFP was added (no inhibition). Assuming the Michaelis-Menten kinetics, the data were fitted by a rescaled function for competitive inhibition:

$$y = (1 - y_{min})\frac{K + [S]}{K(1 + [I]/K_i) + [S]} + y_{min}$$

, where $y$ represents levels of of phosphorylated Cdc25, $y_{min}$ is the background signal obtained from control samples in which no ATP was added, $[S]$ represents the Cdc25 concentration, $K$ indicates the Michaelis-Menten constant for Chk1 phosphorylation of Cdc25, $[I]$ represents the H3-tail concentration, and $K_i$ indicates the inhibition constant. Here, $y$, $y_{min}$, $[S]$, and $[I]$ are experimentally given. To estimate the values of $K$ and $K_i$, we performed global fitting using the two datasets of both $Cdc25^{String}$ and $Cdc25^{Twine}$, where $K_i$ was shared in both datasets while $K$ was independently fitted to each dataset.

### Western blotting

Samples were separated on a 12% TGX Stain-Free FastCast acrylamide gel (Bio-Rad) and transferred to a low fluorescence PVDF membrane. Membranes were incubated overnight in primary antibodies at 4 °C, washed, and incubated in secondary antibodies for 1 hour at room temperature. Mouse anti-H2B antibody (1:5000; Abcam: ab52484), rabbit anti-H3 antibody (1:2000; Abcam: ab1791), rabbit anti-phosphoserine/threonine antibody (1:1000; ECM Biosciences: PP2551), mouse anti-His-Tag antibody (1:1000; Cell Signaling Technology: 2366), rabbit anti-phospho Histone H3 (T11) (1:2000; Abcam ab5168), anti-GFP (1:2000; Millipore-Sigma: G1546), Alexa Fluor 488-conjugated goat anti-mouse IgG antibody (1:5000; Invitrogen: A-11001), Alexa Fluor 546-conjugated goat anti-mouse IgG antibody (1:2000; Invitrogen: A-11003), and Alexa Fluor 647-conjugated goat anti-rabbit IgG antibody (1:5000; A-21244) were used. Fluorescence was detected using a gel imager (Bio-Rad ChemiDoc MP) and quantified in Fiji/ImageJ [9].

### Mathematical modeling

<u>Model description</u>

Based on previous models[10,11], we constructed a model with two ordinary differential equations that drive the embryonic cell cycle oscillation:

$$\frac{d}{dt}Cyc = k_{synth} - k^*_{deg}Cyc \qquad [1]$$

$$\frac{d}{dt}Cyc\text{-}Cdk1 = k_{synth} - k^*_{Wee1}Cyc\text{-}Cdk1 + k^*_{Cdc25}(Cyc - Cyc\text{-}Cdk1) - k^*_{[illegible]}\,Cyc\text{-}Cdk1 \qquad [2]$$

where $Cyc$ represents mitotic cyclins and $Cyc\text{-}Cdk1$ indicates an active form of cyclin-Cdk1 complex. Cyclin synthesis occurs at a constant rate of $k_{synth}$ and the newly-synthesized cyclins rapidly associate with a large pool of Cdk1 and form $Cyc\text{-}Cdk1$. Cyclins are degraded by the first order kinetics with an apparent rate constant of $k^*_{deg}$:

$$k^*_{deg} = a_{deg} + b_{deg}\frac{Cyc\text{-}Cdk1^{n_{deg}}}{K_{deg}^{n_{deg}} + Cyc\text{-}Cdk1^{n_{deg}}} \qquad [3]$$

, which is dependent on cyclin-Cdk1 through cyclin-Cdk1-mediated APC/C activation. The active form of cyclin-Cdk1 can be converted to an inactive state through inhibitory phosphorylation by Wee1 and the inhibitory phosphorylation can be removed by Cdc25. Note that levels of inactive cyclin-Cdk1 (phosphorylated Cyc-Cdk1 at inhibitory phosphorylation sites) are given by $Cyc - Cyc\text{-}Cdk1$ because of mass conservation. Wee1 and Cdc25 activities are also regulated by cyclin-Cdk1, creating double-negative and positive feedback loops that result in ultrasensitive responses of Wee1 and Cdc25 to cyclin-Cdk1, as described by:

$$k^*_{Wee1} = a_{Wee1} + b_{Wee1} \frac{K_{Wee1}^{n_{Wee1}}}{K_{Wee1}^{n_{Wee1}} + Cyc\text{-}Cdk1^{n_{Wee1}}} \quad [4]$$

$$k^*_{Cdc25} = a_{Cdc25} + b_{Cdc25} \frac{Cyc\text{-}Cdk1^{n_{Cdc25}}}{K_{Cdc25}^{n_{Cdc25}} + Cyc\text{-}Cdk1^{n_{Cdc25}}} \quad [5]$$

. The model shows a stable oscillation with appropriate parameter values (discussed below).

Next, we introduced Chk1 into the model. It is well established that Chk1 inhibits cyclin-Cdk1 activity by positively regulating Wee1 and negatively regulating Cdc25[12,13]. Therefore, we rewrite $k^*_{Wee1}$ and $k^*_{Cdc25}$ as follows:

$$k^*_{Wee1} = \left( a_{Wee1} + b_{Wee1} \frac{K_{Wee1}^{n_{Wee1}}}{K_{Wee1}^{n_{Wee1}} + Cyc\text{-}Cdk1^{n_{Wee1}}} \right) (1 + \beta Chk1^*) \quad [4']$$

$$k^*_{Cdc25} = \left( a_{Cdc25} + b_{Cdc25} \frac{Cyc\text{-}Cdk1^{n_{Cdc25}}}{K_{Cdc25}^{n_{Cdc25}} + Cyc\text{-}Cdk1^{n_{Cdc25}}} \right) (1 - Chk1^*) \quad [5']$$

, where we assumed that Chk1 activity, which is represented by $Chk1^*$, linearly affects Wee1 and Cdc25 activities[7]. Note that the equations [4'] and [5'] are equivalent to the equations [4] and [5] in the absence of Chk1 activity (i.e., *chk1* mutant). The Chk1 activity is controlled by the basal Chk1 activity and the concentration of H3 that inhibits Chk1:

$$Chk1^* = Chk1 \frac{1}{1 + H3/K_{i,H3}} \quad [6]$$

, where $Chk1$ is a parameter describing the basal Chk1 activity ($0 \leq Chk1 \leq 1$), $H3$ indicates the H3 concentration, and $K_{i,H3}$ denotes the inhibition constant. When we simulate the effect of H3T11A mutation that results in a weaker inhibitor activity on the Chk1 activity, we rewrite the above equation as:

$$Chk1^* = Chk1 \frac{1}{1 + H3/K_{i,H3} + H3T11A/K_{i,H3T11A}} \quad [7]$$

, where $H3T11A$ represents the concentration of H3T11A, and $K_{i,H3T11A}$ indicates the inhibition constant for H3T11A.

The ordinary differential equations were numerically computed using the scipy.integrate.ode [14].

<u>Parameters</u>

The values of parameters used for simulation are summarized in Tables S2 and S3. Values of parameters involved in the cyclin-Cdk1 processes were essentially the same as the values used in the previous model[11], except: 1) all of the rate constants were rescaled and $b_{deg}$was modified to take into account the difference in the time scale of embryogenesis between *Xenopus* that the previous work modeled and *Drosophila* that we studied in this paper. 2) $k_{synth}$ was slightly decreased with each cycle to recapitulate the increase in cell cycle durations that is independent of Chk1 activity (Fig. S7). Values of parameters involved in Chk1 inhibition by H3 were experimentally estimated (Fig. 2, C and D, and the next section). The basal Chk1 activity, $Chk1$, and $\beta$ were chosen to fit the data.

Phase plane analysis

The equations [1] and [2] define the motion of solutions in the two-dimensional phase plane ($Cyc$ and $Cyc\text{-}Cdk1$). Nullclines divide the phase plane into regions of different qualitative behaviors. The $Cyc$ nullcline is a set of points where $\frac{d}{dt}Cyc = 0$ and is given from equation [1] by:

$$Cyc = \frac{k_{synth}}{k^{*}_{deg}} \qquad [8]$$

, where note that $k^{*}_{deg}$ is a function of $Cyc\text{-}Cdk1$ (equation [3]). Similarly, The $Cyc\text{-}Cdk1$ nullcline that satisfies $\frac{d}{dt}Cyc\text{-}Cdk1 = 0$ is given from equation [2] by:

$$Cyc = \frac{-k_{synth} + \left(k^{*}_{Wee1} + k^{*}_{Cdc25} + k^{*}_{deg}\right)Cyc\text{-}Cdk1}{k^{*}_{Cdc25}} \qquad [9]$$

, where note that $k^{*}_{deg}$, $k^{*}_{Wee1}$, and $k^{*}_{Cdc25}$ are a function of $Cyc\text{-}Cdk1$ (equations [3], [4'], and [5']). The nullclines are plotted in Fig. 4B in blue and red, respectively. Intersection(s) of nullclines is either a stable or unstable fixed point. When Chk1 activity is relatively low, the fixed point is unstable and all trajectories converge to a limit cycle (Fig. 4B, left). By contrast, when Chk1 activity is sufficiently high, the fixed point is stable and the oscillatory behavior is lost (Fig. 4B, right). Thus, increasing Chk1 activity ultimately alters the behavior of the cell cycle and results in cell cycle arrest. Chk1 inhibition by H3 contributes to suppressing premature cell cycle arrest in the wild-type condition but is insufficient when a large amount of H3 is depleted (i.e., SLBP RNAi), resulting in premature cell cycle arrest (Fig. S8).

**Estimation of nuclear H3 concentrations in the early embryo**

We sought to estimate the nuclear H3 concentrations in the early embryo to compare the *in vivo* condition and the $K_i$ value that we obtained in our *in vitro* experiments. Note that this estimation is not exact and a more precise estimation will require more direct and sensitive measurements.

We have previously measured the dynamics of nuclear volumes and relative total nuclear H3 intensities in NC11–13[15]. To estimate absolute H3 concentrations we assumed that total nuclear H3 signal in late telophase/early interphase is approximately equivalent to two genomes worth H3 since DNA replication and H3 import would not have proceeded yet. The genome size of *Drosophila* is approximately 175 Mb[16]. Assuming that nucleosomes are spaced every 200 bps, two genomes worth H3 is $1.75 \times 10^8 \,/\, 200 \times 2 \times 2 = 3.5 \times 10^6$ copies. We are aware that the centromeric histone CID instead of H3 is deposited at centromeres, but the sizes of centromeres are between 101 and 171 Kb[17], which occupies only a tiny fraction of the entire genome. We also note that we did not discriminate between replication-coupled H3 and replication-independent H3.3 in this estimation. Using these values, we estimated that the total nuclear H3 concentration is more than 100 μM at the onset of nuclear cycles due to the small, compact nuclear size and gradually decreases as nuclear volumes grow and chromatin decondenses (Fig. S2). Also, nuclear H3 concentrations decrease with each cycle because non-DNA-bound H3 becomes depleted by the increasing number of nuclei in late cycles [15]. The H3 concentrations at half of the interphase duration (3 min into NC11, 4.5 min into NC12, and 8 min into NC13) were 87 μM, 72 μM, and 52 μM, in NC11, 12, and 13, respectively, and were used as input for simulation. These estimations revealed that the nuclear H3 concentration is comparable to the *Ki* value (47 μM) that we obtained in our *in vitro* kinase assays (Fig. 2, C and D), suggesting that Chk1 inhibition by H3 may be possible at physiological concentrations.

### Supplementary Discussion

#### Other mechanisms for cell cycle slowing independent of Chk1

Although Chk1 activity is crucial for proper cell cycle slowing at the MBT, other Chk1-independent mechanisms also certainly contribute to the regulation of the pre-MBT cell cycles as *chk1* mutant embryo still exhibits small but significant increases in cell cycle durations before the MBT. This is also evidenced by the requirement for Chk1 and histone-independent modulation of the apparent cyclin synthesis in our model. Dilution of DNA replication factors or cyclins, early zygotic transcription of cell cycle inhibitors such as Tribbles and Frs, the conflict of transcription and replication forks, and dNTP availability appear to affect the cell cycle in the early embryo[6,18–23]. Additionally, time-dependent mechanisms play a role in maternal RNA clearance that coincides with the MBT. It is not currently feasible to determine the relative effects of these factors on the pre-MBT cell cycle as they are tightly dependent on each other and the system is highly nonlinear. A major challenge to the field will be to incorporate all of these elements into a single framework.

#### The Chk1 inhibition in contexts other than the pre-MBT cell cycles

The predominant regulatory role of histones and histone posttranslational modifications as transcriptional regulators through their essential function as chromatin components cannot be overstated. However, the focus on the function of histones in the context of chromatin may have caused us to overlook the regulatory role(s) for histones outside of the nucleosome. Future studies will be required to address how frequently extra-chromosomal histones act as signaling molecules and if their functions extend past the regulation of Chk1. The mechanism of Chk1 inhibition described here may continue beyond the pre-MBT embryo into other developmental and disease contexts. For example, histone null embryos fail to accumulate $Cdc25^{String}$ in the first zygotically controlled embryonic cell cycle[24], suggesting that H3 inhibition of Chk1 may be a more general feature of cell cycle regulation. Histone levels are often abnormal in rapidly dividing cancer cells which also frequently have defects in DNA damage checkpoint response[25]. Interestingly, a mutant for Rad53, a functional analog of Chk1, results in the accumulation of excess histones in yeast, suggesting that excess histones may further enhance their own accumulation[26]. Moreover, Chk1 need not be the sole target for regulation by nucleoplasmic or cytoplasmic histone pools. Substrate competition has been implicated to play a significant role in the regulation of the cell cycle[27,28]. In theory, any protein that interacts with histone tails on chromatin could also be regulated by excess histone off chromatin.

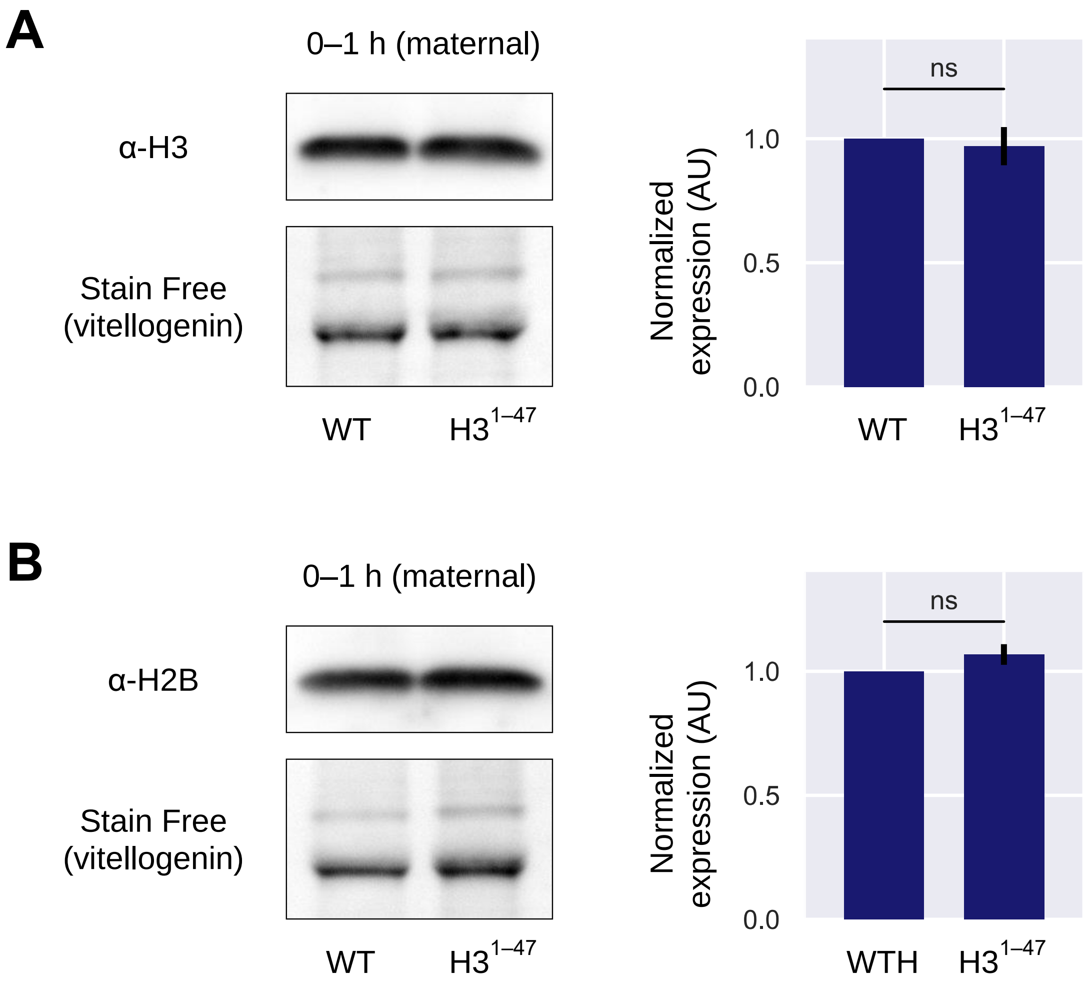


**Fig. S1. Expression of H3-tail does not affect total histone levels.**

(A) and (B) Levels of maternally deposited H3 and H2B protein in embryos produced from WT females or matα4-tubGal4>UASz-H31-47-mEGFP females. Embryos were collected over a one-hour period and the lysates were analyzed by western blotting. H3 and H2B signals were normalized to total protein using vitellogenin in the Stain-Free channel. Histone levels were unaffected by H3-tail expression. Data represent the mean ± SEM of three biological replicates.

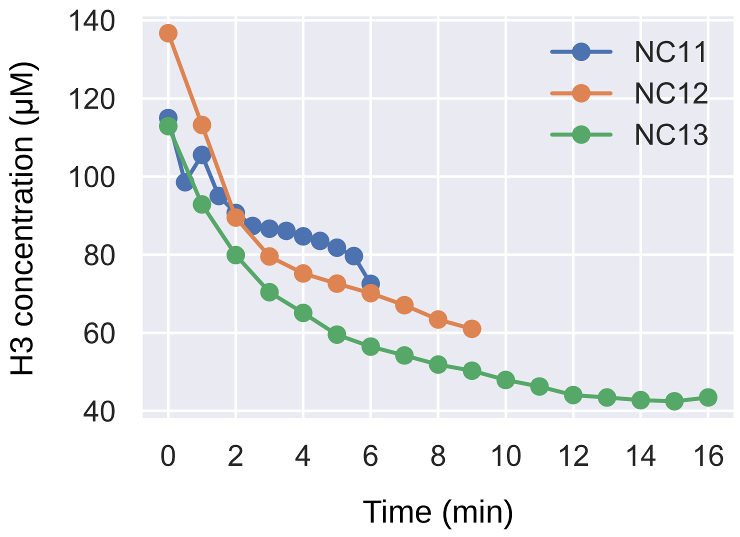


**Fig. S2. Estimation of nuclear H3 concentrations.**

Estimated concentrations of nuclear H3 in the interphase nucleus are shown. We have previously measured nuclear volumes and relative levels of nuclear H3 in NC11–13[15]. Using the fluorescence intensities of H3 in late telophase/early interphase that would correspond to two genomes worth histones, we estimated nuclear H3 concentrations in the early embryo (see Materials and Methods for details).

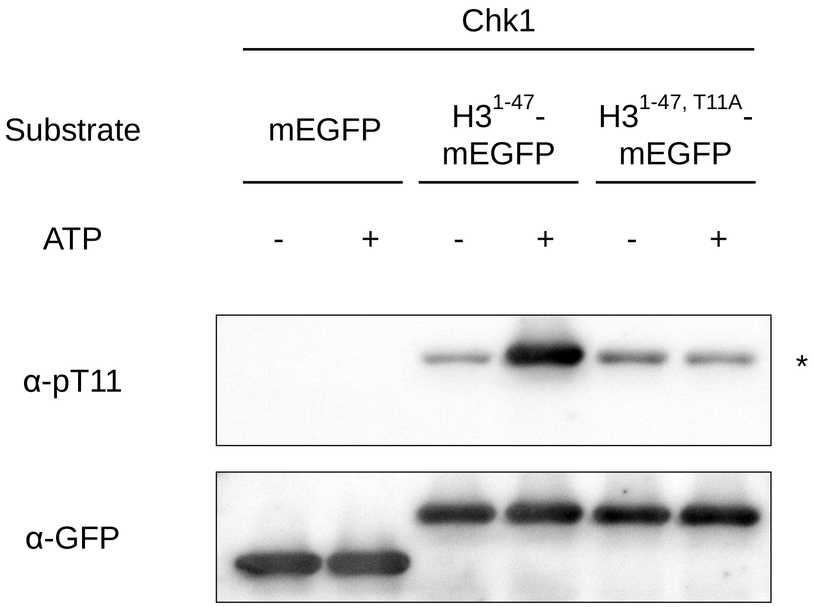


**Fig. S3. Chk1 phosphorylates Thr 11 in the H3-tail in vitro.**

0.06 μM of Chk1 was incubated with 2 μM of mEGFP, $H3^{1\text{-}47}$-mEGFP, or $H3^{1\text{-}47,\ T11A}$-mEGFP and levels of phosphorylated Thr 11 were detected by western blotting. Note that a weak nonspecific signal was present in the absence of the Thr 11 residue and is denoted as an asterisk.

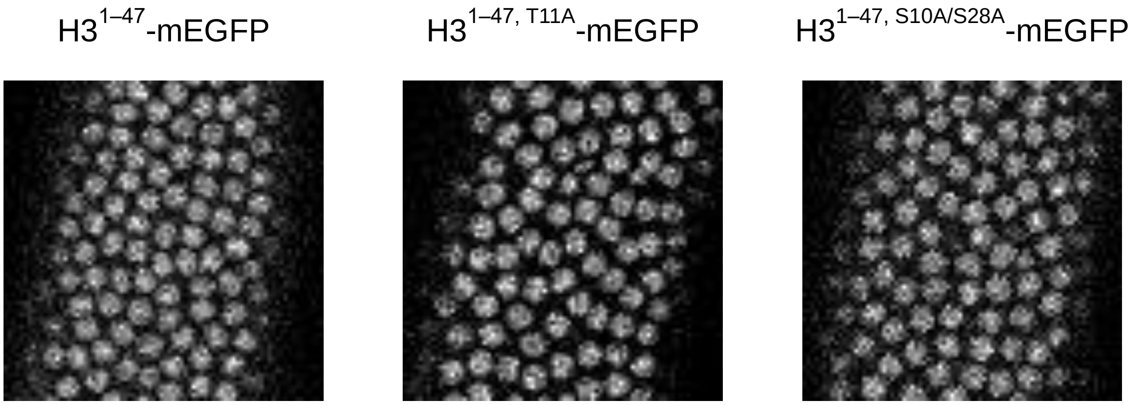


**Fig. S4. H3-tail mutants do not affect localization.**

Confocal images of H3$^{1-47}$-mEGFP, H3$^{1-47, T11A}$-mEGFP, and H3$^{1-47, S10A/S28A}$-mEGFP at NC13 are shown. Localization patterns of H3-tail were unaffected by T11A or S10A/28A mutation.

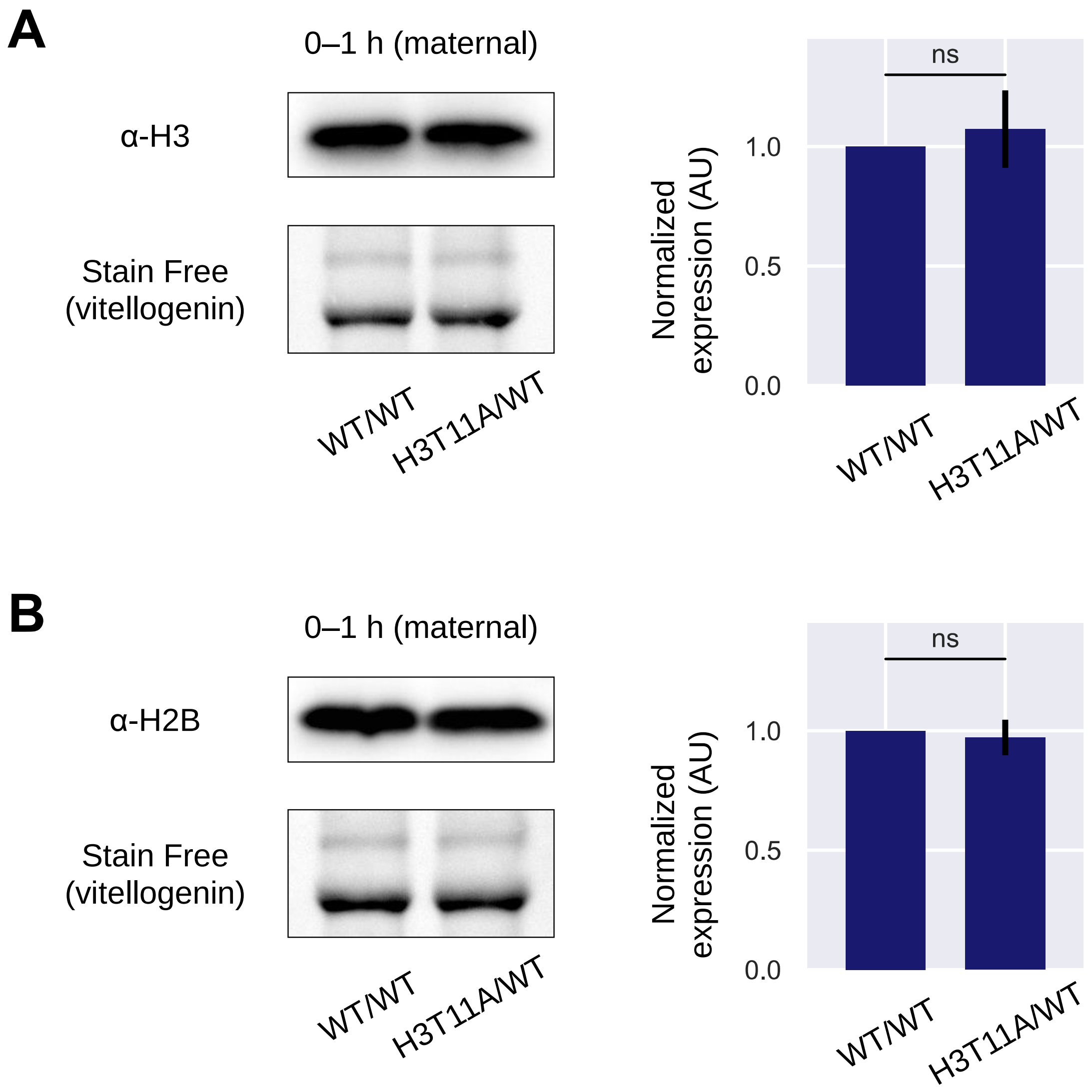


**Fig. S5. Heterozygous H3T11A mutation in the engineered histone gene cluster does not affect total histone levels.**

Levels of maternally deposited H2B and H3 protein in embryos produced from females homozygous for WT His-GUs (10X His-GUs$^{WT}$/10X His-GUs$^{WT}$) and heterozygous for H3T11A (10X His-GUs$^{H3T11A}$/10X His-GUs$^{WT}$). Data represent the mean ± SEM of three biological replicates. Note that previous work has shown that total embryonic histone mRNA and protein levels are unaffected by the reduced copy number[2,29,30] consistent with our observation that the WT His-GUs embryos have cell cycle times comparable to true WT embryos.

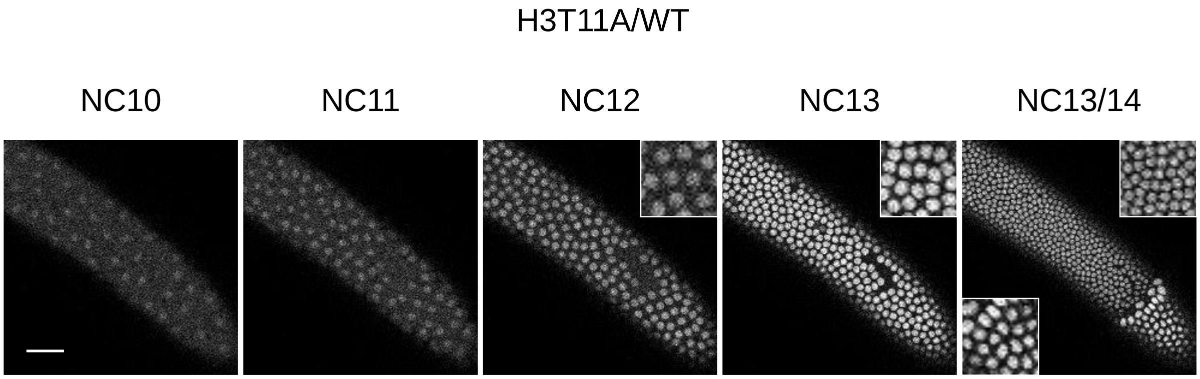


**Fig. S6. Embryo heterozygous for H3T11A undergoes partial premature cell cycle slowing.**

Confocal images of NLS-mRFP in an embryo heterozygous for H3T11A, in which some nuclei stopped dividing at NC13. Insets, enlarged images. Scale bar, 50 μm.

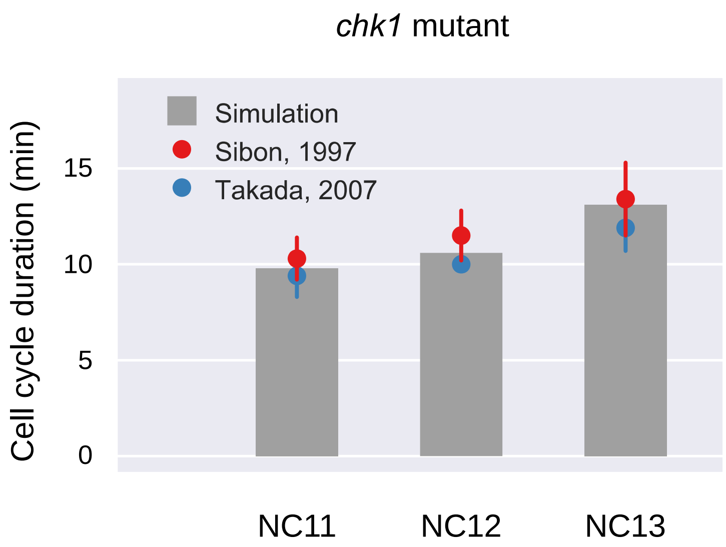


**Fig. S7. Simulated cell cycle durations in chk1 mutant embryos.**

Although embryos produced from homozygous *chk1* females have fast cell cycles due to the lack of Chk1 activity, there is still a significant increase in cell cycle durations in NC11–13[31,32]. To account for the Chk1-independent cell cycle slowing, we slightly decreased the apparent rate of basal cyclin synthesis with each cycle, recapitulating the cell cycle dynamics in the absence of Chk1 activity.

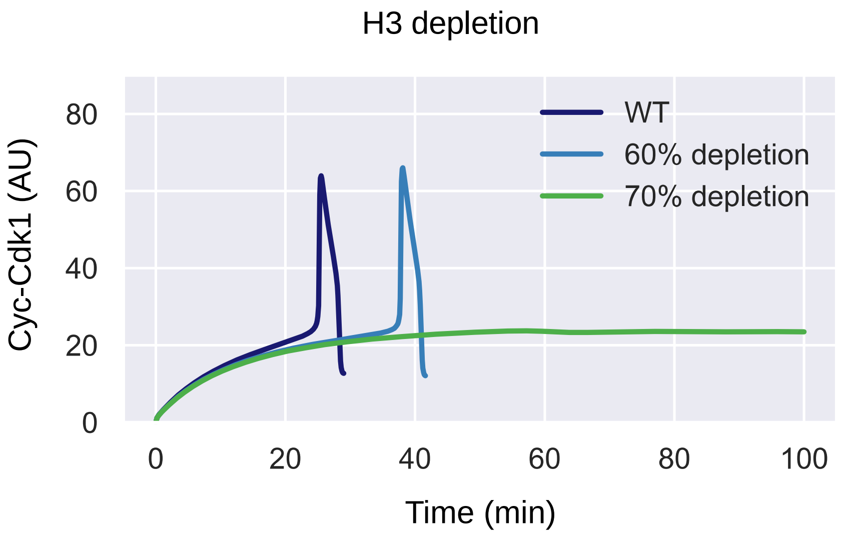


**Fig. S8. Simulation recapitulates premature cell cycle arrest in H3 depleted embryos.**

Simulated dynamics of Cyc-Cdk1 in NC13 with respect to different levels of H3 depletion are shown. H3 depletion up to 69% results in prolonged but stable cell cycle oscillations while more than 70% depletion results in cell cycle arrest that is represented by a single stable steady state.

**Table S1. The number of nuclear divisions.**

| | | Cell cycle paused at | | |
|---|---|---|---|---|
| **Genotype** | **n** | **NC13** | **NC14** | **NC15** |
| WT | 43 | 0% (0) | 100% (43) | 0% (0) |
| $H3^{1\text{-}47}$-mEGFP | 16 | 0% (0) | 81% (13) | 19% (3) |
| 10X His-$GUs^{WT}$ / 10X His-$GUs^{WT}$ | 17 | 0% (0) | 100% (17) | 0% (0) |
| 10X His-$GUs^{H3T11A}$ / 10X His-$GUs^{WT}$ | 14 | 14% (2) | 86% (12) | 0% (0) |

**Table S2. Parameters values used for simulation.**

| **Parameter** | **Value**[*] |
|---|---|
| $k_{synth}$ | 0.0112 (NC11), 0.0105 (NC12), 0.0095 (NC13) |
| $a_{deg}$ | 0.112 |
| $b_{deg}$ | 0.224 |
| $K_{deg}$ | 0.032 |
| $n_{deg}$ | 17 |
| $a_{Cdc25}$ | 1.792 |
| $b_{Cdc25}$ | 8.96 |
| $K_{Cdc25}$ | 0.035 |
| $n_{Cdc25}$ | 11 |
| $a_{Wee1}$ | 0.896 |
| $b_{Wee1}$ | 4.48 |
| $K_{Wee1}$ | 0.030 |
| $n_{Wee1}$ | 3.5 |
| $\beta$ | 0.7 |
| $Chk1$ | 0.2 |
| $K_{i,H3}$ | 47 [μM] |
| $K_{i,H3T11A}$ | 159 [μM] |

[*]All units are shown in terms of an arbitrary unit of concentration and minute except where otherwise explicitly indicated.

**Table S3. Values of H3 concentrations used for simulation.**

| Genotype | H3 [μM] | H3T11A [μM] |
|---|---|---|
| WT (NC11) | 87 | 0 |
| WT (NC12) | 72 | 0 |
| WT (NC13) | 52 | 0 |
| H3-tail | 52 × 1.5 | 0 |
| H3T11A-tail | 52 | 52 × 0.5 |
| H3T11A/WT | 52 × 0.6 | 52 × 0.4 |
| H3 depletion (60% depletion) | 52 × 0.4 | 0 |
| H3 depletion (70% depletion) | 52 × 0.3 | 0 |
